# Molecularly distinct models of zebrafish *Myc*-induced B cell leukemia

**DOI:** 10.1101/446245

**Authors:** Chiara Borga, Clay A. Foster, Sowmya Iyer, Sara P. Garcia, David M. Langenau, J. Kimble Frazer

## Abstract

Zebrafish models of T cell acute lymphoblastic leukemia (T-ALL) have been studied for over a decade, but curiously, robust zebrafish B cell ALL (B-ALL) models had not been described. Recently, our laboratories reported two seemingly closely-related models of zebrafish B-ALL. In these genetic lines, the primary difference is expression of either murine or human transgenic c-MYC, each controlled by the zebrafish *rag2* promoter. Here, we compare ALL gene expression in both models. Surprisingly, we find that B-ALL arise in different B cell lineages, with *ighm*^+^ vs. *ighz*^+^ B-ALL driven by murine *Myc* vs. human *MYC*, respectively. Moreover, these B-ALL types exhibit signatures of distinct molecular pathways, further unexpected dissimilarity. Thus, despite sharing analogous genetic makeup, the ALL types in each model are markedly different, proving subtle genetic changes can profoundly impact model organism phenotypes. Investigating the mechanistic differences between mouse and human c-MYC in these contexts may reveal key functional aspects governing MYC-driven oncogenesis in human malignancies.

---

Zebrafish are a valuable leukemia model due to highly conserved hematopoietic and oncogenic pathways, facile genetics, and ease of use in chemical genetic screens. However, until recently, robust zebrafish B-cell leukemia models had not been described^1, 2^. The first transgenic zebrafish leukemia model was created 15 years ago and targeted murine c-*Myc* (*mMyc*) to thymocytes of AB strain zebrafish, leading to the rapid development of T-cell acute lymphoblastic leukemia (T-ALL)^3^. Additional genetic models were subsequently developed that result in induction of T-ALL^4^, but B-cell leukemia models lagged behind^5, 6^.

In two recent reports published in *Leukemia*, our groups independently demonstrated the development of zebrafish B-ALL using transgenic expression of *mMyc* or human c-*MYC* (*hMYC*) controlled by the zebrafish recombination activating gene 2 (*rag2*) promoter^1, 2^. Both models shared genetic and phenotypic features, but there were also key differences including strain background and species differences in the MYC transgene that was used to generate each model. Here, we compare and contrast these models, making the important finding that zebrafish develop at least four molecularly distinct ALL types, including cortical thymocyte-arrested *cd4^+^*/*cd8^+^* T-ALL, *ighm*^+^ B-ALL, *ighz*^+^ B-ALL, and biphenotypic T/B-ALL.

Thirteen ALLs were purified from *rag2:hMYC;lck:eGFP* double-transgenic fish^2^. As previously reported, these leukemias had heterogeneous GFP expression, with T-ALL being exclusively GFP^hi^, B-ALL exclusively GFP^lo^, and other fish harboring mixed-ALL with both GFP^hi^ and GFP^lo^ cells, representing simultaneous T- and B-ALL, respectively^2^. Leukemias were subjected to RNA-seq transcriptomic profiling and compared with transplanted leukemias generated from single ALL clones described by Garcia et al., 2018^1^. Principal Component Analysis clearly distinguished *mMyc-*induced T-ALL from B-ALL, with the single *mMyc-* induced biphenotypic B/T-ALL clustering between these samples (Fig. 1A). The eight *hMYC*-induced ALLs that were largely GFP^hi^ clustered with known T-ALLs (2, 6, 8-11, 13, 14), while three primarily GFP^lo^ ALL clustered near the *mMyc* B-ALLs (3-5). Two *hMYC*-induced ALLs with substantial populations of both GFP^lo^ and GFP^hi^ cells grouped near the *mMyc* biphenotypic B/T leukemia. Hierarchical clustering using the top 100 positively- and negatively-correlated genes from PC2 confirmed that these genes defined B and T lymphocytes, respectively (Fig. 1B), with B-ALLs expressing *cd79b*, *syk*, *pax5*, *blnk* and *efb1*, while T-ALLs expressed *cd8b*, *lck*, *runx3* and *gata3*. As previously reported, the biphenotypic B/T-ALL and mixed *hMYC*-induced ALLs expressed both T- and B-cell lineage genes^1, 2^. PC2 up-regulated genes were enriched for B cell signaling pathways when independently assessed by GSEAsig (Supplemental Tables 1 and 2).

**Figure 1.**
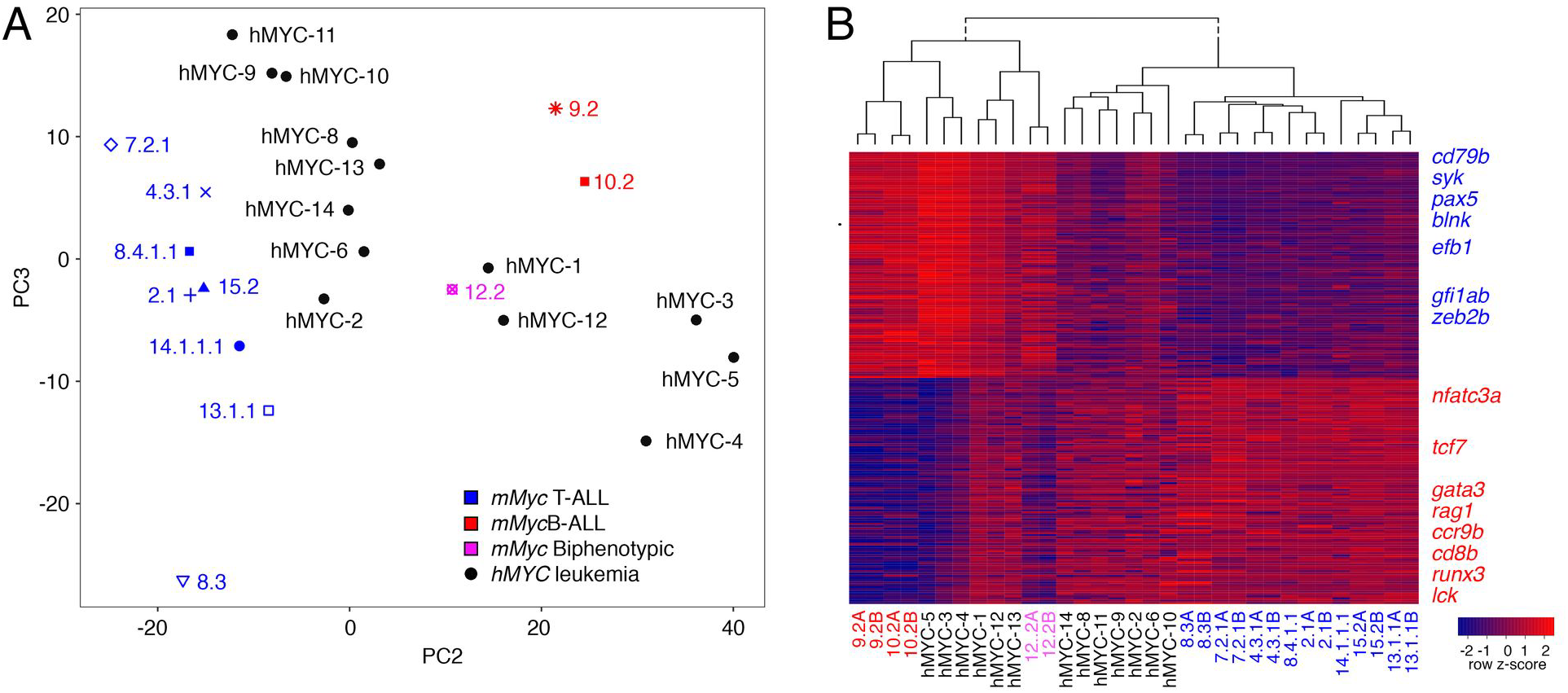
Gene expression differences define T-vs. B-lineages in ALL of both the *rag2:mMyc* and *rag2:hMYC;lck:eGFP* models. (**A**) Principal component analysis of RNAseq expression profiles of previously-classified *mMyc-*induced T-(blue, n=8), B-(red, n=2), or biphenotypic ALL (pink, n=1) and compared with 13 unknown *hMYC-*induced ALL (black). (**B**) Heat map and hierarchical clustering using the top 100 positively- and negatively-correlated genes from PC2. B cell-specific genes are denoted in blue and T cell-specific genes in red at the right.

To examine clonality and maturation in *mMyc-* and *hMYC-*induced ALL, we next analyzed the expression of constant and variable regions of the T cell receptor β (*tcrβ* and immunoglobulins μ and ζ (*ighm*, *ighz*; Fig. 2A). Every *mMyc* and *hMYC* T-ALL exhibited *tcrβ* expression, with V(D)J recombination occurring in most samples as determined by expression of specific variable regions^1^. Conversely, *mMyc* and *hMYC* B-ALL did not recombine or express *tcrβ*, but expressed constant regions of *ighm* or *ighz*. Ig variable regions were not detected, indicating V(D)J rearrangement likely had not occurred in these leukemias and suggesting B-ALLs arrest at the early pro-B cell stage. As expected, *hMYC* mixed-ALL contained distinct T- and B-ALL clones expressing both *tcr* and *ig* mRNAs, with their relative expression correlating well with the percentage of GFP^hi^/T-ALL vs. GFP^lo^/B-ALL cells found in each sample (Fig. 2A). Intriguingly, *mMyc* B-ALL expressed exclusively *ighm* while *hMYC* B-ALL favored *ighz* expression, indicating that *mMyc* and *hMYC* might be oncogenic in distinct B cell lineages.

**Figure 2.**
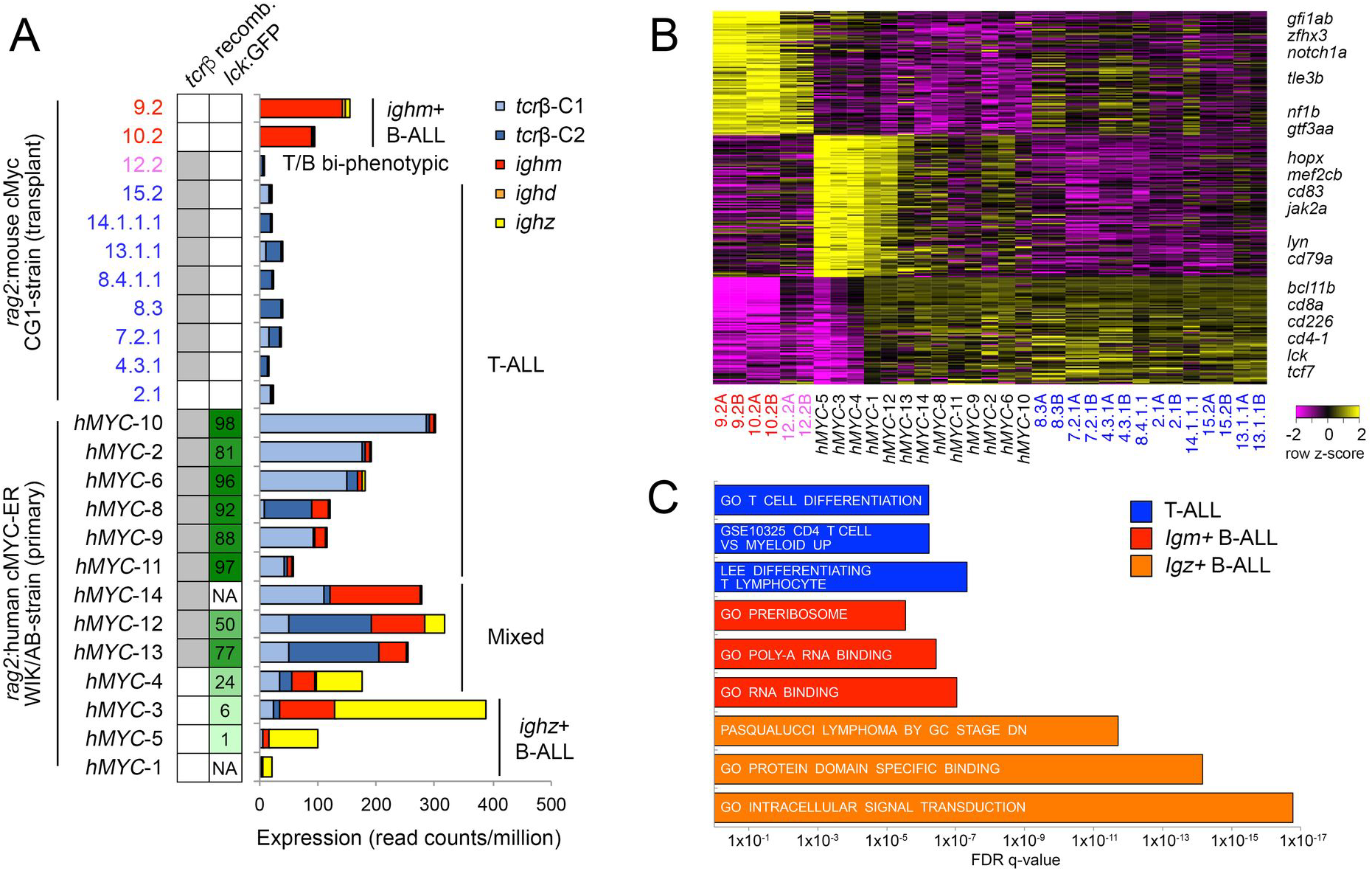
Identification of two molecularly-distinct B-ALL types arising independently in either the *ighm^+^* or *ighz^+^* B cell lineages. (**A**) T cell receptor beta and Ig heavy chain expression in individual ALLs. *tcrβ* recombination is denoted by grey-shaded boxes (left column), with percentage of GFP^hi^ cells in each *rag2:hMYC;lck:eGFP* ALL noted in right column. Histograms depict expression of *tcrβ* and *igh* constant regions by each ALL. Not available (NA). (**B**) Heatmap showing expression of genes differentially expressed in T-ALL, *mMyc*/*ighm*^+^ B-ALL, and *hMYC*/*ighz*^+^ B-ALL. (**C**) Gene set enrichment analysis using genes positively correlated with each ALL molecular subtype. T-ALL (blue), *mMyc*/*ighm*^+^ B-ALL (red), and *hMYC*/*ighz*^+^ B-ALL (orange). Complete geneset and GSEAsig results are provided in Supp. Tables 3 and 4.

To further explore differences between these models, we next identified genes uniquely-expressed by T-ALL, *mMyc*/*ighm*^+^ B-ALL, or *hMYC*/*ighz*^+^ B-ALL (Fig. 2B). As expected T-ALLs expressed known T cell lineage markers, yet *mMyc*/*ighm^+^* and *hMYC*/*ighz*^+^ B-ALLs were transcriptionally distinct. *mMyc*/*ighm*^+^ B-ALL expressed *gfi1ab, zfhx3, notch1a, nf1b*, and *gtf3aa*. By contrast, *hMYC*/*ighz*^+^ B-ALL expressed higher *cd79a*, *cd83, mef2cb*, and *jak2a* levels. To further test for differences in these two molecular subtypes of B-ALL, we next performed GSEAsig using these same differentially-regulated genes. From this analysis, we uncovered that *mMyc*/*ighm*^+^ B-ALLs exhibited significant enrichment for pathways regulating ribosome biogenesis and RNA binding (Fig. 2C and Supplemental Tables 3 and 4). By contrast, *hMYC*/*ighz*^+^ B-ALLs were enriched for intracellular signaling, protein binding, and germinal center B cell maturation pathways. In support of our findings, Liu et al. recently reported the identification of molecularly and biological distinct *ighz*^+^ and *ighm*^+^ B cell lineages using *rag2:mCherry; cd79b:GFP* transgenic zebrafish^7^. In the context of normal B cell development, *ighz^+^* B cells were mCherry^hi^/GFP^lo^ while *ighm*^+^ B cells were mCherry^hi^/GFP^hi^. Overall, these results demonstrate that *mMyc*/*ighm*^+^ and *hMYC*/*ighz*^+^ B-ALLs are not subtle B cell leukemia variants, but rather distinct malignancies that arise in different B cell types with vastly different molecular pathway signatures.

In summary, although zebrafish B cell leukemia models were lacking for many years, our analyses reveal two highly-divergent types of B-ALL. This is surprising, as both models utilize the same promoter (*rag2*) to regulate a near-identical oncoprotein, c-Myc/MYC, with the only differences being the MYC transgene species of origin and the genetic backgrounds upon which the models were developed. Yet, despite the high molecular similarity of both models, these B-ALL subtypes also show unique gene expression signatures when compared to one another, which likely reflects differences in both their lineage (*ighm* vs. *ighz*) and potential differences in MYC transcriptional targets expressed by the early developmental stages of these distinct pro-B cell populations. Our new analysis of these models reconciles the perceived differences in the manuscripts published by our groups, identifying four molecularly distinct ALL subtypes in zebrafish: cortical *cd4^+^*/*cd8^+^* T-ALL, biphenotypic B/T ALL, *ighm*^+^ B-ALL, and *ighz*^+^ B-ALL. Developing a wider array of leukemia models and refining mechanisms that drive their growth, aggression, and stem cell frequency will surely lead to new insights into human disease.

